# Cell type directed design of synthetic enhancers

**DOI:** 10.1101/2022.07.26.501466

**Authors:** Ibrahim Ihsan Taskiran, Katina I. Spanier, Valerie Christiaens, David Mauduit, Stein Aerts

## Abstract

Transcriptional enhancers act as docking stations for combinations of transcription factors and thereby regulate spatiotemporal activation of their target genes. A single enhancer, of a few hundred base pairs in length, can autonomously and independently of its location and orientation drive cell-type specific expression of a gene or transgene. It has been a long-standing goal in the field to decode the regulatory logic of an enhancer and to understand the details of how spatiotemporal gene expression is encoded in an enhancer sequence. Recently, deep learning models have yielded unprecedented insight into the enhancer code, and well-trained models are reaching a level of understanding that may be close to complete. As a consequence, we hypothesized that deep learning models can be used to guide the directed design of synthetic, cell type specific enhancers, and that this process would allow for a detailed tracing of all enhancer features at nucleotide-level resolution. Here we implemented and compared three different design strategies, each built on a deep learning model: (1) directed sequence evolution; (2) directed iterative motif implanting; and (3) generative design. We evaluated the function of fully synthetic enhancers to specifically target Kenyon cells in the fruit fly brain using transgenic animals. We then exploited this concept further by creating “dual-code” enhancers that target two cell types, and minimal enhancers smaller than 50 base pairs that are fully functional. By examining the trajectories followed during state space searches towards functional enhancers, we could accurately define the enhancer code as the optimal strength, combination, and relative distance of TF activator motifs, and the absence of TF repressor motifs. Finally, we applied the same three strategies to successfully design human enhancers. In conclusion, enhancer design guided by deep learning leads to better understanding of how enhancers work and shows that their code can be exploited to manipulate cell states.

## Introduction

A long-standing goal in genome biology is to decipher the regulatory code of transcriptional enhancers and to build predictive models of gene regulation. The current paradigm of how enhancers yield spatiotemporal gene expression is that they act as binding platforms for specific combinations of transcription factors (TF), that can be activators, repressors, or neutral/pioneering TFs. Cooperative or hierarchical binding of multiple TFs results in nucleosome eviction at the enhancer, followed by cofactor recruitment and interaction with other enhancers, with the proximal promoter, and with RNA polymerase, through which the rate of transcription initiation is modulated ^1–4^. Cell type specific expression of a target gene is achieved when a unique combination of TFs activates a specific enhancer; while this enhancer remains either passively (“default-off” ^5,6^) or actively repressed in other cell types (e.g., via repressor binding ^7^ or co-repressor/polycomb recruitment). Typically, when an enhancer is translocated to another chromosome or to an episomal plasmid, it maintains cell type specific control of its nearby reporter gene ^8,9^. Therefore, its regulatory capacity is contained within the enhancer DNA sequence, and has co-evolved to respond uniquely to a specific trans-environment in a cell type. A thorough understanding of how enhancer activation is encoded in its DNA sequence is important, as it is a key component for the modelling and prediction of gene expression ^10^; for the interpretation of non-coding genome variation ^11,12^; and for the reconstruction of dynamic gene regulatory networks underlying developmental, homeostatic, and disease-related cell states.

Many complementary approaches and techniques have been used to decode enhancer logic ^8^. These include studies of individual enhancers by mutational analysis ^13^, in vitro TF binding (e.g., electrophoresis mobility shift assay), cross-species conservation ^14^, and reporter assays. When such studies were scaled up ^15–17^, common features of co-regulated enhancers could be identified. These experimental findings also triggered the improvement of computational methods for the prediction of *cis*-regulatory modules, whereby feature selection and parameter optimization led to new insights into how binding sites cluster and how their strength (or binding energy) impacts enhancer function ^18–23^. Wider adoption of genome-wide profiling of chromatin accessibility ^24^, single-cell chromatin accessibility ^25–27^, histone modifications ^28,29^, TF binding ^30^, and enhancer activity ^16,31^ led to significantly larger training sets of co-regulated enhancers that could then be used for *a posteriori* discoveries of TF motifs and enhancer rules, aided by the growing resources of high-quality TF motifs ^32,33^. Additional mechanistic insight has been provided by thermodynamic modelling of enhancers ^34,35^, in vivo imaging of enhancer activity ^36^, the analysis of genetic variation through eQTL and caQTL analysis ^6,37^, and high-throughput in vitro binding assays ^38,39^. Recently, the enhancer biology field embraced the use of convolutional neural networks (CNN) and network-explainability techniques that again provided a significant leap forward in terms of prediction accuracy and syntax formulation ^10,40–44^.

An orthogonal strategy to decode enhancer logic is to engineer synthetic enhancers from scratch. This approach has the advantage that the designer knows exactly which features are implanted, so that the minimal requirements for enhancer function can be revealed. Recent work showed the promise of CNN-driven enhancer design by successfully designing yeast promoters ^45^, and by using a CNN to select high-scoring enhancers for S2 cells, from a large pool of random sequences ^42^. Here we tackle the next challenge in enhancer design, namely to design enhancers that are cell type specific. To this end, we implement and compare three different CNN-based strategies to design enhancers, including the use of Generative Adversarial Networks (GAN) ^46^ (Fig. 1). We further use these strategies to remove target cell types from, or add new target cell types to existing enhancers. These techniques also allow designing minimal enhancers and to evolve near-enhancer regions from the genome into functional enhancers. Finally, we computationally evaluate the different design processes to draw new insight into animal enhancer codes.

**Figure 1:**
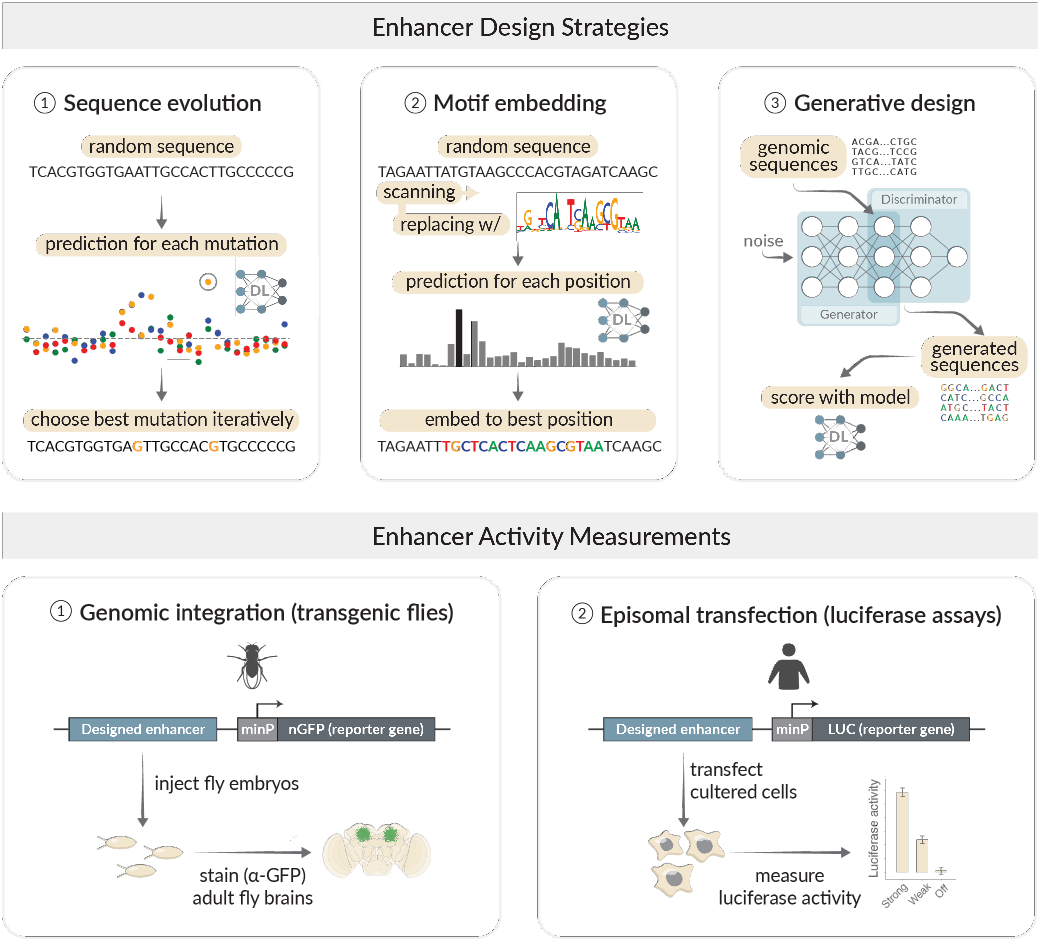
Deep learning-based enhancer design. Overview of three enhancer design strategies and activity measurements of designed enhancers in Drosophila brains (via site directed integration in the genome) and human cell lines (via transfection). Each of the design strategies makes use of Deep Learning (DL) models that were previously trained and validated on cell type specific chromatin accessibility data, namely DeepFlyBrain ^43^ and DeepMEL^44^.

## Results

### Enhancer design by in silico evolution

As a first strategy for enhancer design, we created synthetic enhancers to specifically target Kenyon cells (KC) in the mushroom body of the fruit fly brain, using a nucleotide-by-nucleotide sequence evolution approach (Methods). This approach starts from a 500 bp random sequence that is evolved *in silico* towards a chosen cell type through multiple iterations, using the DeepFlyBrain ^43^ model to calculate fitness scores. At each iteration we performed saturation mutagenesis whereby all nucleotides were mutated one by one, and each sequence variation was scored by DeepFlyBrain to select the mutation with the greatest positive delta score for the KC class (Fig. 2a). We performed this procedure starting from 6000 random sequences and observed that after 15 iterations, DeepFlyBrain prediction scores increased from around the minimal score (0) to nearly the maximum score (1) (Fig. 2b).

**Figure 2:**
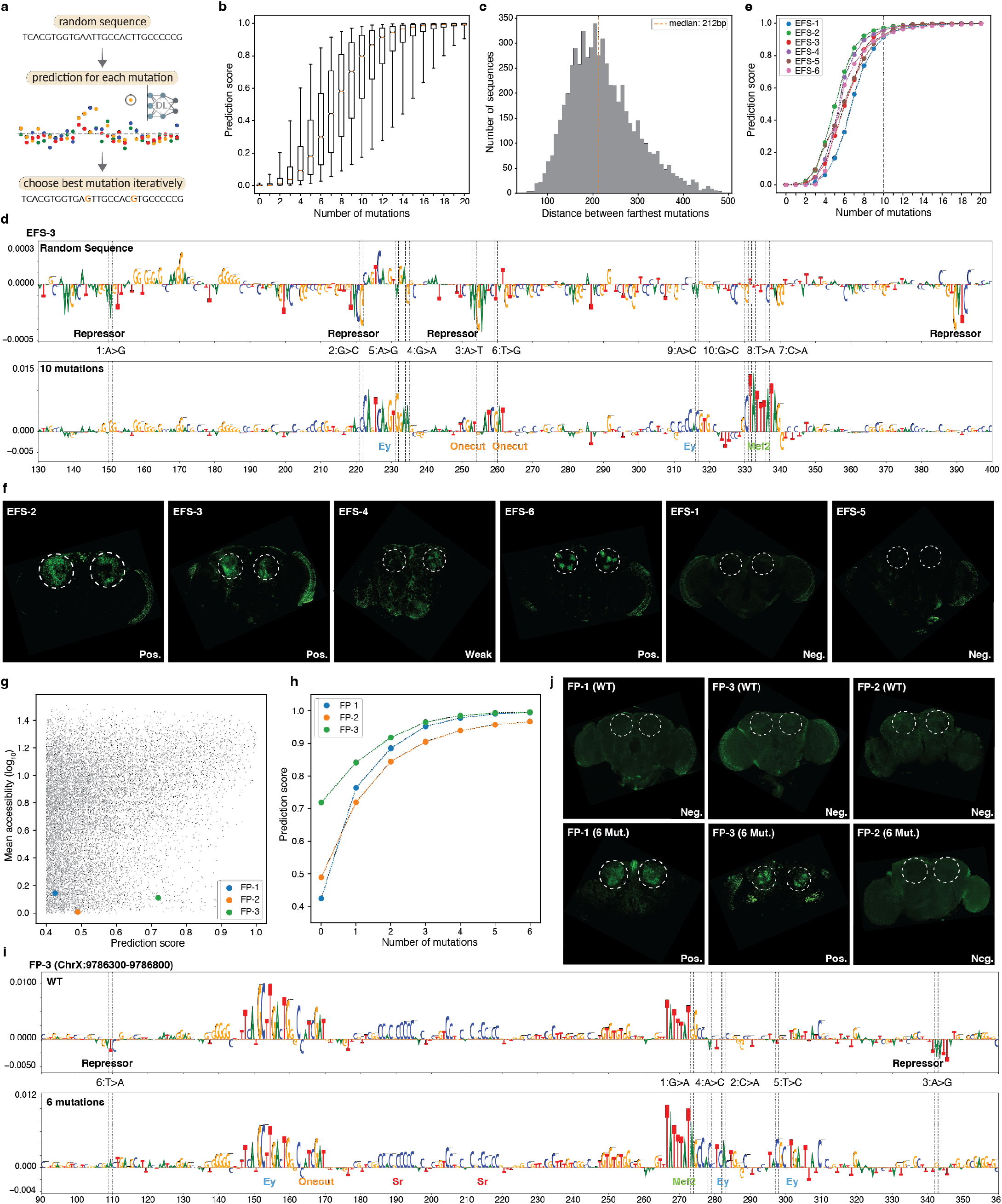
In silico sequence evolution towards functional enhancers. **a**, In silico sequence evolution approach. **b**, Prediction score distribution of the sequences for the γ-KC class (*n* = 6,000) after each mutation. The box plots show the median (center line), interquartile range (box limits), and 5th and 95th percentile range (whiskers). **c**, Distribution of distances (*n* = 6,000) between farthest mutations on each sequence after 10 iterative mutations. The orange line shows the median. **d**, Nucleotide contribution scores of a selected random sequence in its initial form (top) and after 10 iterations (bottom). The position of the mutations are shown with dashed lines. The mutational order is written in between top and bottom plots together with the type of nucleotide substitutions. **e**, Prediction scores of six selected sequences at each mutational step. The selected iteration (10^th^ mutation) is indicated with a dashed line. **f**, In vivo enhancer activity of the cloned sequences with 10 mutations. The expected location of KC is shown with dashed circles. **g**, Comparison between γ-KC prediction score and mean γ-KC accessibility for the binned fly genome regions. The selected regions with high prediction and low accessibility are highlighted with blue, orange, and green dots. **h**, Prediction scores of the selected sequences for each mutational step. **i**, Nucleotide contribution scores of a selected genomic sequence in its initial form (top) and after 6 iterations (bottom). The position of the mutations are shown with dashed lines. The mutational order is written in between top and bottom plots together with the type of nucleotide substitutions. **j**, In vivo enhancer activity of the cloned genomic sequences with 6 mutations. The expected location of KC is shown with dashed circles.

Next, we investigated the effect of the initial (random) sequence, and the specific paths that are followed through the search space towards local optima. For only a small fraction (3%) of random sequences, a high prediction score was reached at later iterations, and the prediction score remained below 0.5 even after 15 mutations (Supplementary Fig. 1a). These sequences were mostly characterized by more instances of repressor binding sites together with an increased number of mutations required to generate main activator binding sites. A second observation is that even though 500 bp space is given to the model, the selected mutations accumulated in about 200 bp space (Fig. 2c).

Next, we investigated the consequences of each mutation on shaping the enhancer code using DeepExplainer-based contribution scores ^47,48^ (Methods) (Fig. 2d). This revealed that initial random sequences usually harbor several short repressor binding sites by chance and these are preferentially destroyed during the first iterations. These repressor sites contribute negatively to the KC class and represent candidate binding sites for KC specific repressor TFs such as Mamo and CAATTA ^49^. Interestingly, the nucleotides with the highest impact represent mutations that destroy a repressor binding site and simultaneously generate a binding site for the key activators Eyeless, Mef2 or Onecut (Fig. 2d, Supplementary Fig. 1b). Eventually, at the local optima, DeepExplainer revealed multiple activator binding sites, whereby Ey, Mef2, and Onecut sites dominate (Fig. 2d, Supplementary Fig. 1b).

To test whether the in silico evolved enhancers can drive reporter gene expression in vivo, we randomly selected six sequences after 10 iterations (Fig. 2d,e, Supplementary Fig. 1b,c) and tested them in vivo with a GFP reporter in a transgenic fly line (Methods). Four out of these six tested synthetic enhancers were active specifically in the targeted cell-type, the Kenyon cells (Fig. 2f). This strategy thus represents an intuitive and efficient approach to generate cell type specific enhancers.

Given that KC enhancers can arise from random sequences after ∼10 mutations, we hypothesized that certain genomic regions may require even fewer mutations to acquire KC enhancer activity. We scanned the entire fly genome and identified regions with high prediction scores but without chromatin accessibility in KC (Fig. 2g, Methods). By applying sequence evolution to these sequences with only 6 mutations (Fig. 2h,i, Supplementary Fig. 2a), we successfully utilized their potential, yielding 2 out of 3 positive KC enhancers (Fig. 2j). This suggests that KC enhancers, and likely other cell type enhancers as well, can arise de novo in the genome with few mutations.

### Enhancer design towards multiple cell type codes

A single enhancer can be active in multiple, different cell types ^50^, and our earlier work suggested that this can be achieved by enhancers that contain multiple codes for different cell types, intertwined within a single ∼500 bp sequence ^43^. Based on this finding, we wondered whether a genomic enhancer that is active in a single cell type, could be synthetically augmented to become also active in a second cell type.

To test this, we started from the previously characterized *CG15117* enhancer, that is accessible and active (Fig. 3a,b) in T1 neurons ^43^. Its activity pattern is also predicted correctly by DeepFlyBrain (Fig. 3c). We then performed in silico evolution on this enhancer towards the KC cell type, similarly as done above. However, at each iteration, we now choose the mutation causing the strongest increase for the KC prediction score while simultaneously maintaining a high T1 prediction score (Fig. 3d, Methods). After 14 mutations, the T1 enhancer was also predicted as KC enhancer (Fig. 3d). The nucleotide contribution scores before and after the in silico evolution show that the T1 enhancer code was barely touched, with the main binding sites for Awh and Oc (TFs) still present (Fig. 3e, top). On the other hand, all repressor binding sites for the KC code were destroyed and typical KC activator binding sites for Ey and Sr were created (Fig. 3e, bottom). Testing the initial and augmented sequence in vivo with a GFP reporter confirmed the spatial expansion of the enhancer activity to a second cell type (Fig. 3b,f). We repeated this procedure for another optic lobe enhancer, again adding KC activity, which required 13 mutations (Fig. 3g-k, Supplementary Fig. 3a).

**Figure 3:**
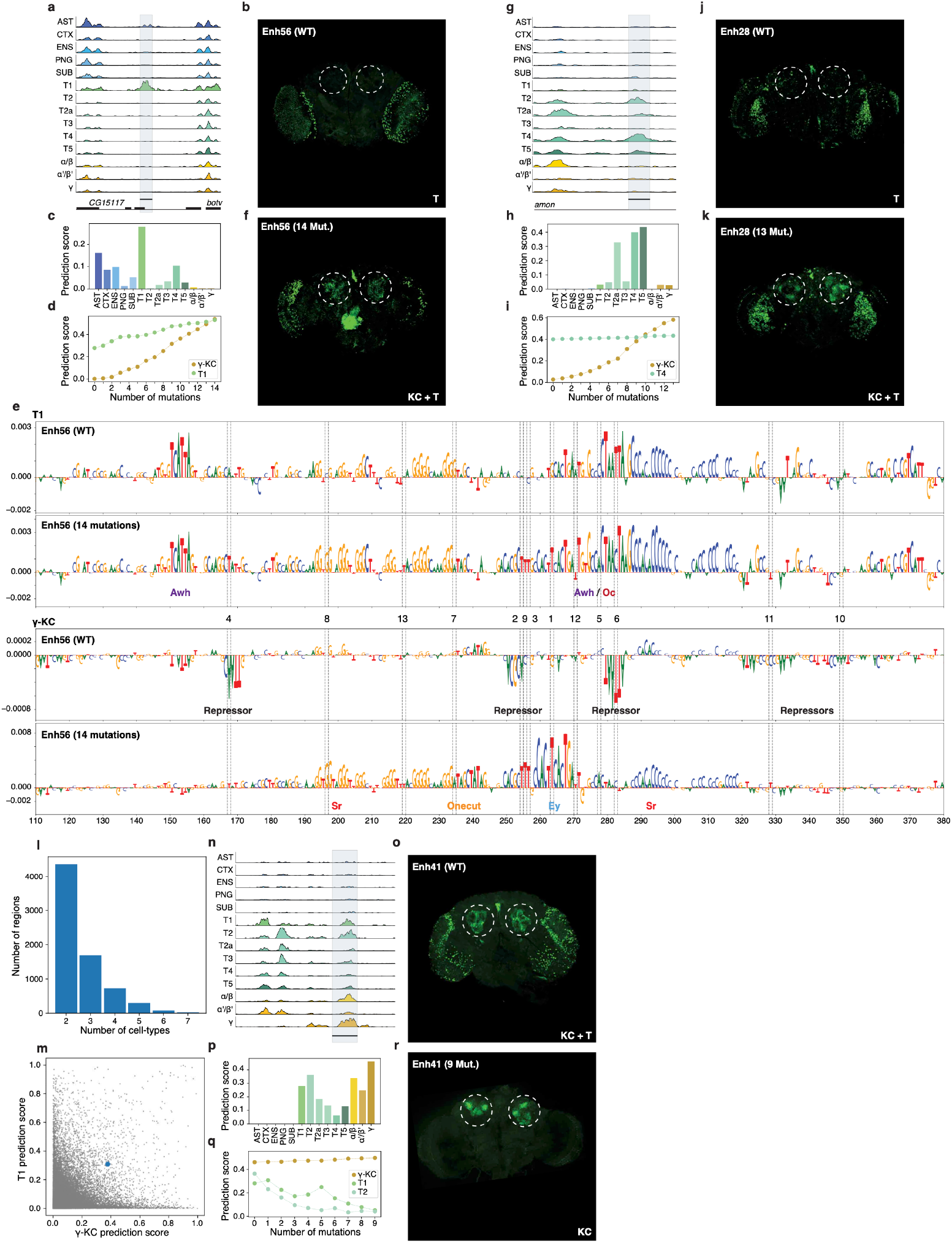
Spatial expansion and restriction of enhancer activity. **a**, Chromatin accessibility profile near *CG15117* gene. **b**, In vivo enhancer activity of the *CG15117* enhancer. **c**, *CG15117* enhancer prediction scores for each cell-type. **d**, Prediction scores for the γ-KC and T1 classes after each mutational step. **e**, Nucleotide contribution scores of wild-type (WT) sequence and after 14 mutations for T1 (top) and γ-KC (bottom). The position of the mutations are shown with dashed lines. The mutational order is written in-between top and bottom plots. **f**, In vivo enhancer activity of the *CG15117* enhancer after 14 mutations. The expected location of KC is shown with dashed circles. **g**, Chromatin accessibility profile near the *amon* gene. **h**, *amon* enhancer prediction scores for each cell-type. **i**, Prediction scores for the γ-KC and T4 classes after each mutational step. **j**, In vivo enhancer activity of the *amon* enhancer. **k**, In vivo enhancer activity of the *amon* enhancer after 13 mutations. The expected location of KC is shown with dashed circles. **l**, Number of regions that score high (>0.3) for multiple cell-types. **m**, Comparison between γ-KC prediction score and T1 prediction score for the accessible regions in fly brain (*n* = 95,931). The selected region with high γ-KC and T1 prediction is highlighted with a blue dot. **n**, Chromatin accessibility profile of this region in multiple cell-types. **o**, In vivo enhancer activity of the *Pkc53e* enhancer. **p**, *Pkc53e* enhancer prediction scores for each cell-type. **q**, Prediction scores for the γ-KC, T1, and T2 classes after each mutational step. **r**, In vivo enhancer activity of the multi cell-type enhancer after 9 mutations. The expected location of KC is shown with dashed circles.

These results also suggest that, reciprocally, enhancers active in multiple cell types may be pruned towards a single cell-type code. We searched for genomic enhancers that score high for multiple cell-types (Fig. 3l). We selected a *Pkc53e* enhancer (Fig. 3m) that is accessible and active (Fig. 3n,o) in both optic lobe T-neurons and KCs and predicted correctly by the model (Fig. 3p). This time, we drove the in silico evolution to maintain the KC prediction score, while decreasing the T-neurons prediction score (Methods). After 9 mutations, the sequence was predicted to have only KC activity (Fig. 3q). Nucleotide contribution scores show that the most important binding sites for KCs were unaffected after 9 mutations, while the activator binding sites were destroyed and new repressor binding sites were created for T-neurons (Supplementary Fig. 3b). Testing the initial and the final sequence in vivo with a GFP reporter confirmed the spatial restriction of the enhancer activity (Fig. 3o,r). Together, our results suggest that intertwining enhancer codes can be independently dissected and altered, guided by the DeepFlyBrain model.

### Enhancer design by motif implantation

As a second strategy, we used a motif implantation approach to design KC enhancers. We started from a random sequence and in each iteration, we implanted a strong motif instance for a certain TF at every possible position, then scored each sequence with DeepFlyBrain, and selected the location with the highest gain towards the gamma-KC class. The rationale behind this strategy came from our results above, where nucleotide-by-nucleotide sequence evolution showed that all the selected mutations were associated with the creation or destruction of a transcription factor binding site, rather than affecting contextual sequence between motif instances (Fig. 2d,i, 3e, Supplementary Fig. 1b, 2a, 3a,b). This suggested that a combination of appropriately positioned activator motifs, without the presence of repressor motifs, would be sufficient to create a cell type specific enhancer in the fly brain. Furthermore, we reasoned that by applying this design strategy to thousands of random sequences, and each time selecting high-scoring architectures, we could gain additional insight into the KC enhancer logic. To this end, we created 2000 random sequences and first implanted a single binding site for one of the four key activators of KC enhancers, namely Ey, Mef2, Onecut, and Sr ^49^, and then specific combinations of sites in a particular implantation order (Methods). This revealed that Ey and Mef2 had the strongest effect on the prediction score, while One-cut and Sr increased the prediction score only marginally (Fig. 4b). Implanting Ey and Mef2 consecutively increased the score more than the sum of their individual contribution and their implantation order did not affect the final score (Fig. 4b). Adding Onecut and then Sr on top of Ey and Mef2 sites increased the scores even further until it reached the level that we obtained above after 15 mutations through in silico sequence evolution (Fig. 4b). We also found that high-scoring configurations consisted of activator sites that are positioned close together within a distance usually smaller than 100 bp (Fig. 4c,d, Supplementary Fig. 4a). When the Ey and Mef2 pair were implanted on the same strand, we observed strong preference for a 5 bp distance between the two binding sites whereby Mef2 was located upstream of Ey (Fig. 4c, Supplementary Fig. 4b). This distance was reduced to 4 bp when Ey and Mef2 sites were implanted on opposite strands of each other (Supplementary Fig. 4b). For the Ey and Onecut pair, distances in the range of 100 bp were equally preferred, however there was a strong preference for a 3 bp space and Onecut preferred the downstream side of Ey (Fig. 4d, Supplementary Fig. 4c).

**Figure 4:**
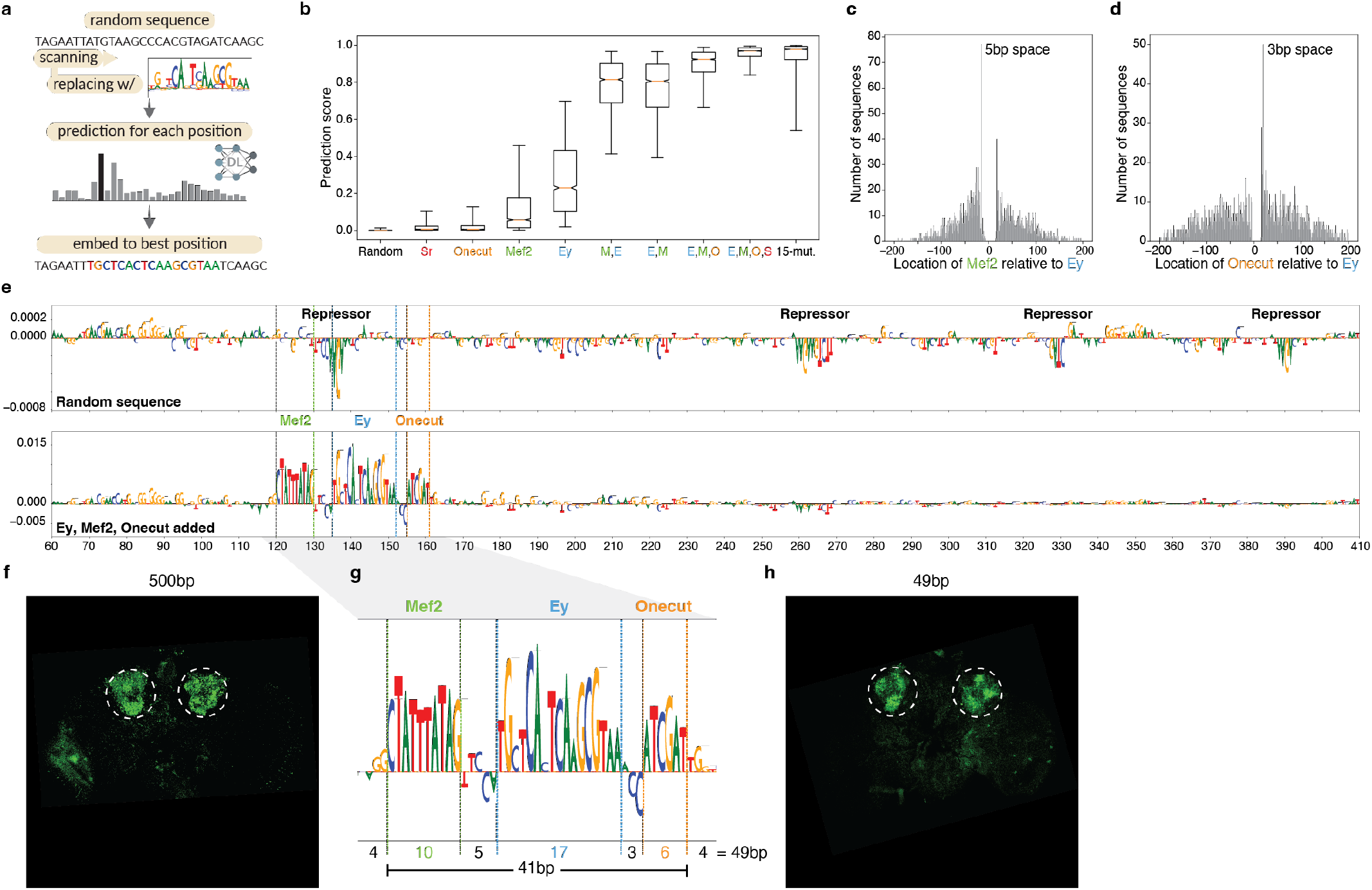
Motif implantation towards minimal enhancer design. **a**, Motif implantation approach. **b**, Prediction score distribution of the sequences for the γ-KC class (*n* = 2,000) after each motif implantation and after 15 mutations. Abbreviations are used for Ey (E), Mef2 (M), Onecut (O), and Sr (S). The box plots show the median (center line), interquartile range (box limits), and 5th and 95th percentile range (whiskers). **c**, Distribution of Mef2 locations relative to Ey (*n* = 2,000). **d**, Distribution of Onecut locations relative to Ey (*n* = 2,000). **e**, Nucleotide contribution scores of a selected random sequence in its initial form (top) and after Ey, Mef2, and Onecut implantations (bottom). The position of the motifs are shown with dashed lines. **f**, In vivo enhancer activity of the cloned 500 bp sequence with Ey, Mef2, and Onecut implantations. The expected location of KC is shown with dashed circles. **g**, Zoom into the selected 49 bp part of the 500 bp sequence from **e**. The size of the motifs, the spaces in between motifs, and the flankings are shown at the bottom. **h**, In vivo enhancer activity of the cloned 49 bp sequence with Ey, Mef2, and Onecut implantations. The expected location of γ-KC is shown with dashed circles.

Next, we calculated nucleotide contribution scores before and after motif implantations of an example sequence where motifs were inserted close together (Fig. 4e). The initial random sequence contained multiple repressor binding sites and the Ey binding site implantation destroyed one of the repressor binding sites. Mef2 and Onecut implantations followed the predicted spacing relative to Ey, with a distance of 5 bp and 3 bp, respectively. Even though some repressor binding sites were still present at further distances, their relative negative contribution was decreased after the activator binding site implantations (Fig. 4e). Testing this designed 500 bp sequence in vivo confirmed specific activity in KC (Fig. 4f). Furthermore, a 49 bp subsequence, containing just the three binding sites, resulted in the same activity and specificity in vivo (Fig 4g,h). This result suggests that a functional KC enhancer can be created via motif-by-motif implantation with just these three binding sites and its size can be decreased to the minimal length required to contain these binding sites.

### Enhancer design by Generative Adversarial Networks

As a third strategy for enhancer design, we used Generative Adversarial Networks (GAN) that have been shown to be powerful generators in different fields ^46,51,52^. We trained a GAN on genomic sequences, using KC enhancers as a training set. After training, we used the generator part of the model to generate synthetic sequences (Fig. 5a, Methods). To evaluate the GAN-generated sequences we used the DeepFlyBrain model to compare the generated sequences with randomly generated sequences using a Markov model (Methods). After 20k batch iterations, the GAN model started outperforming a 5th order background model (Fig. 5b). Moreover, the GAN model captured high-order genomic sequence features (e.g. GC content is higher in the center) while the background model failed as it only focuses on k-mer frequencies and fails to capture any location bias (Fig. 5c). Calculating nucleotide contribution scores of the generated sequences revealed relevant binding sites, although these sites were weaker compared to the sequences generated by the first two strategies (Fig. 5d, Supplementary Fig. 5a). We selected 5 of the generated sequences that were predicted as KC and harbored a combination of weak/strong Ey, Mef2, Onecut and Sr binding sites. Testing their activity in vivo showed that the sequences with better binding sites drove clearer GFP expression in KC (Fig. 5e). In conclusion, GAN generated sequences can also produce functional and cell type specific enhancers, but require a posteriori filtering with pre-trained enhancer models.

**Figure 5:**
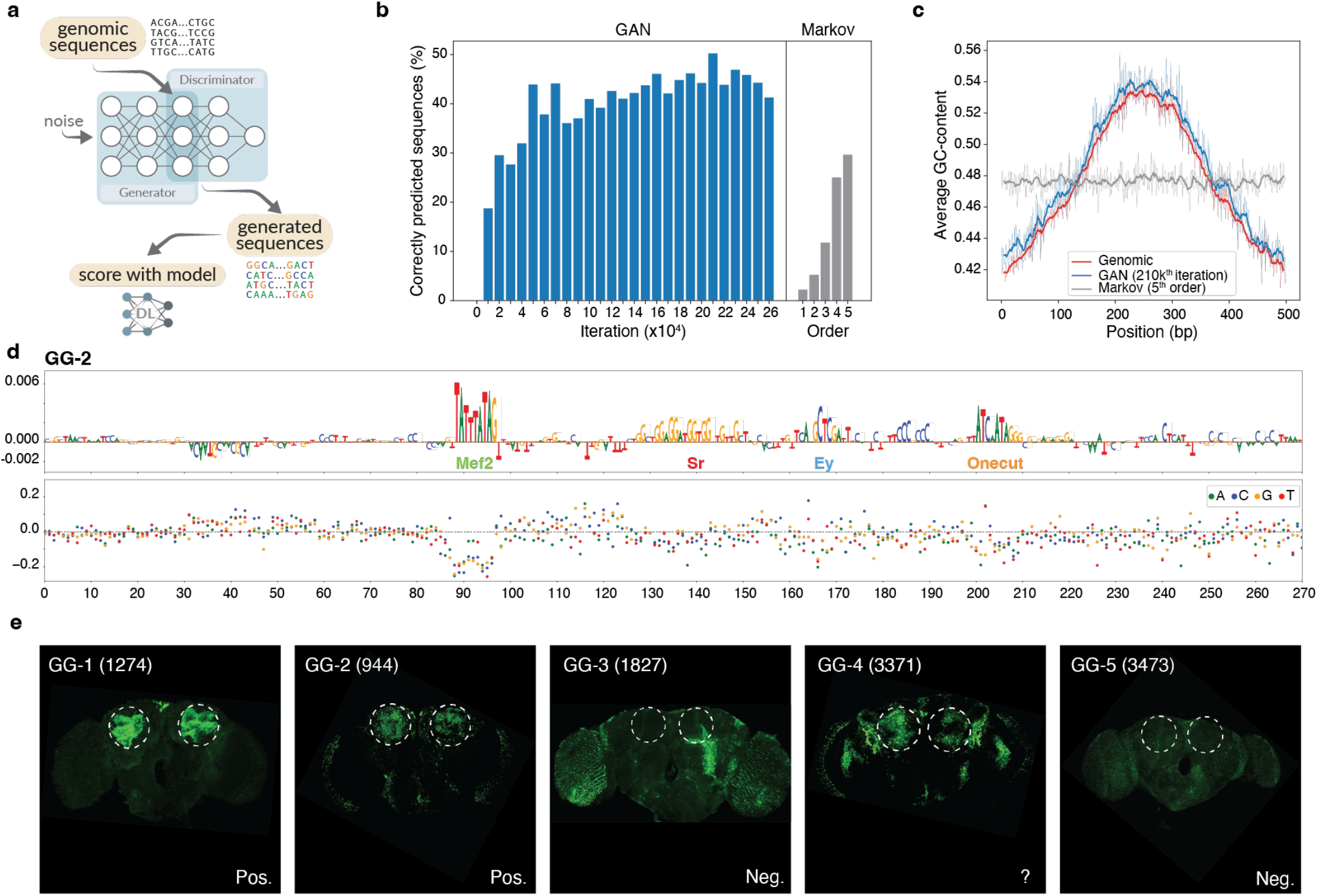
Generative design of enhancers. **a**, Generative design approach. **b**, Percentage of correctly predicted sequences (prediction score > 0.25) that are generated after each 10,000 batch iterations of the GAN model (*n* = 6144 for each iteration) and that are generated by a Markov model with different orders (*n* = 6144 for each order). **c**, Average GC-content over 500 bp genomic (*n* = 6126), GAN-generated at 210k^th^ batch iteration (*n* = 6144), and 5^th^ order Markov-generated sequences (*n* = 6144). **d**, Nucleotide contribution scores of a selected GAN-generated sequence (top) and its in silico saturation mutagenesis assay (bottom). Each dot on the saturation mutagenesis plot represents a single mutation and its effect on the prediction score (*y* axis). **e**, In vivo enhancer activity of the cloned GAN-generated sequences. The expected location of KC is shown with dashed circles.

### Human enhancer design

Next, we investigated whether the same three strategies can also be used to design human enhancers. For this purpose, we designed enhancers for melanocytic (MEL) melanoma cells that we studied previously, and for which we trained and validated two deep learning models, called DeepMEL and DeepMEL2 ^11,44^.

Similar to the *Drosophila* experiments, we started with random sequences and evolved them into human MEL enhancers by following the nucleotide-by-nucleotide sequence evolution approach. The prediction scores started to saturate after 20 mutations (Fig. 6a). We randomly selected 10 regions that were evolved from scratch (EFS1-10) with 15 mutations and tested their activity with a luciferase assay in vitro, in a melanocytic melanoma cell line (MM001) (Methods). 7 out of 10 tested enhancers showed activity in the range of previously characterized positive control (native) enhancers ^53^ and none of them showed activity in a cell line that represents another melanoma cell state (mesenchymal-like, MM047) where the MEL specific TFs (SOX10, MITF, and TFAP2) are not expressed (Fig. 6b, Supplementary Fig. 6a). The nucleotide contribution scores of the positive enhancers showed that the selected mutations destroyed repressor binding sites (e.g. ZEB) and created activator binding sites similar to what we observed in *Drosophila* experiments (Fig. 6d, Supplementary Fig. 6b).

**Figure 6:**
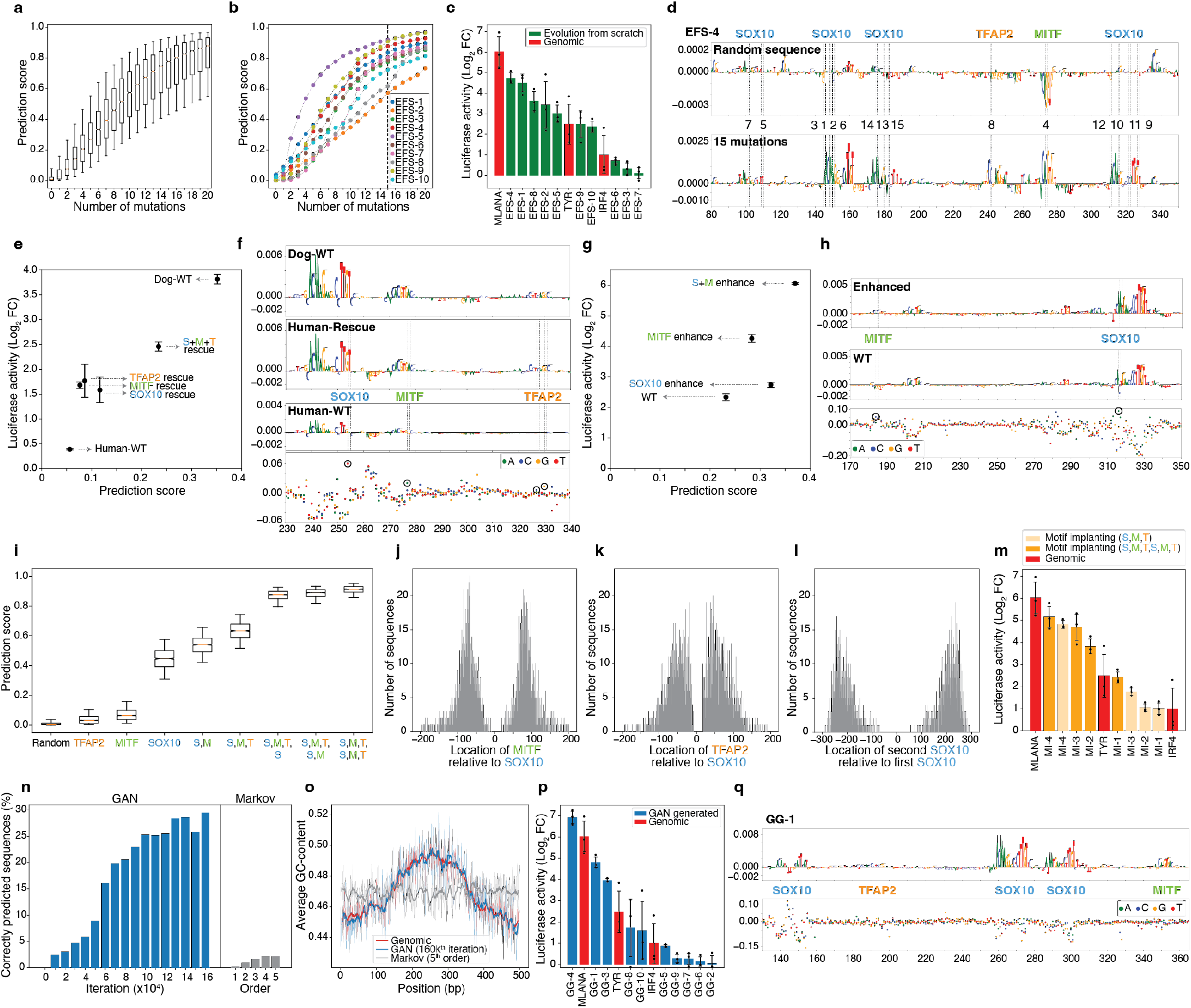
Human enhancer design. **a**, Prediction score distribution of the sequences for the MEL class (*n* = 4000) after each mutation. **b**, Prediction scores of 10 selected sequences after each mutational step. The selected iteration (15^th^ mutation) is indicated with a dashed line. **c**, Bar plot showing the mean luciferase signal (log_2_ fold-change over Renilla) of the synthetic sequences, which were generated by in silico sequence evolution, and genomic enhancers. **d**, Nucleotide contribution scores of a selected synthetic sequence in its initial form (top) and after 15 iterations (bottom). The position of the mutations are shown with dashed lines. The mutational order is written in between top and bottom plots. The name of the generated binding sites by the mutations are indicated at the top. **e**, Dot plot showing the mean luciferase signal (log_2_ fold-change over Renilla) versus prediction score for the MEL class of the wild-type (WT) human and dog genomic sequences and the rescued human sequences. **f**, Nucleotide contribution scores of the dog, human-rescued, and human-WT sequences (top 3 rows) and in silico saturation mutagenesis assay of human-WT sequence (bottom). **g**, Dot plot showing the mean luciferase signal (log_2_ fold-change over Renilla) versus prediction score for the MEL class of the wild-type and enhanced enhancers. **h**, Nucleotide contribution scores of the wild-type (middle) and enhanced (top) enhancers and in silico saturation mutagenesis assay of wild-type enhancer (bottom). **i**, Prediction score distribution of the sequences for the MEL class (*n* = 2,000) after each motif implantation. **j**, Distribution of MITF locations relative to SOX10 (*n* = 2,000). **k**, Distribution of TFAP2 locations relative to SOX10 (*n* = 2,000). **l**, Distribution of the second SOX10 locations relative to the first SOX10 (*n* = 2,000). **m**, Bar plot showing the mean luciferase signal (log_2_ fold-change over Renilla) of the synthetic sequences, which were generated by motif implanting, and genomic enhancers. **n**, Percentage of correctly predicted sequences (prediction score > 0.15) that are generated after each 10,000 batch iteration of the GAN model (*n* = 3968 for each iteration) and that are generated by Markov model with different orders (*n* = 3968 for each order). **o**, Average GC-content over 500 bp genomic (*n* = 3885), GAN-generated at 160k^th^ batch iteration (*n* = 3968), and 5^th^ order Markov-generated sequences (*n* = 3968). **p**, Bar plot showing the mean luciferase signal (log_2_ fold-change over Renilla) of the GAN-generated sequences and genomic enhancers. **q**, Nucleotide contribution scores of a selected GAN-generated sequence (top) and its in silico saturation mutagenesis assay (bottom). The name of the identified binding sites are written in between top and bottom plots. In a, **i**, the box plots show the median (center line), interquartile range (box limits), and 5th and 95th percentile range (whiskers). In **c, e, g, m, p**, the error bars show the standard error of the mean (*n* = 3 biological replicates). In **e, g, i, m**, abbreviations are used for SOX10 (S), MITF (M), and TFAP2 (T). In **f, h, q**, each dot on the saturation mutagenesis plot represents a single mutation and its effect on the prediction score (*y* axis). In **d, f, h**, the position of the mutations are shown with dashed lines and circles.

In the fly brain, we applied in silico sequence evolution to create enhancers from genomic regions with high scores that did not show chromatin accessibility, and could be considered as ‘near-enhancer’ sequences. We also extended this approach to MEL enhancers. We started from a human sequence that has no MEL enhancer activity, but its homologous sequence in the dog genome is accessible and active as MEL enhancer ^44^. We used DeepMEL to introduce 4 mutations that restored the activator binding sites in the human sequence, resulting in a rescue of the activity, as measured by luciferase activity (Fig. 6e,f). As a final variation of this approach, we introduced 2 mutations in a weak MEL enhancer (one to increase the SOX10 binding site affinity and another to convert the ZEB repressor binding site into an MITF activator binding site) which resulted in a 10-fold increase in enhancer activity (Fig. 6g-h).

We also applied the motif implantation strategy to design human enhancers. We implanted SOX10, MITF, and TFAP2 binding sites to 2000 random sequences of 500bp. While implanting only MITF or TFAP2 resulted in a small increase in the prediction score, implanting SOX10 alone had the strongest effect (Fig. 6i). Adding MITF and then TFAP2 on top of SOX10 sites increased the prediction scores to 0.6 on average. The prediction scores continued increasing even further after adding another set of SOX10, MITF, and TFAP2 binding sites (Fig. 6i). We did not observe a single preferential location for the implantation of MITF or TFAP2 relative to SOX10, however both binding sites were located within 100 bp of SOX10 (Fig. 6j,k). The second SOX10 binding sites were placed further away at a 200-250 bp distance relative to the first SOX10 (Fig. 6l). We selected 4 sequences where SOX10, MITF, and TFAP2 were implanted once and also their double-binding-sites version and tested their activity with luciferase assays. All enhancers showed activity in the range of native enhancers ^53^ and adding the binding sites twice always increased the activity of the enhancers (Fig. 6m, Supplementary Fig. 7a,b,c). Replacing implanted binding sites with their weaker versions taken from a native enhancer (IRF4) decreased the activity of the enhancers dramatically (Supplementary Fig. 7a,b,c). To confirm that the activity of the enhancers were driven by the implanted binding sites, we cut the sequences from the most upstream binding site till the most downstream binding site. These subsequences (116-164 bp) were also active with a slight change on their activity levels (Supplementary Fig. 7a,b,c). Finally, instead of choosing the best location for MITF and TFAP2 implantation, we implanted them at the closest location from the SOX10 binding site that would result in a positive change in the prediction score. These shortened sequences (51-64 bp) were as active as their longer (500 bp) version (Supplementary Fig. 7a,b,c).

Finally, we tested the GAN based sequence generation approach in our human model. Very similar to what we observed in the *Drosophila* case, the GAN model started out-performing background models after 20k batch iterations (Fig. 6n) and again captured the high-order sequence features much better than a Markov model (Fig. 6o). We selected 10 GAN-generated sequences (GG1-10) from the best batch iteration that contain SOX10, MITF, and TFAP2 binding sites (Fig. 6p,q, Supplementary Fig. 8a). Testing them in vitro with a luciferase assay showed that 5 of them were active in the range of genomic enhancers ^53^ (Fig. 6p). Observed activity levels in a mesenchymal-like (MES) melanoma line were overall higher (meaning, their cell type specificity is lower) for the GAN-generate enhancers, compared to the enhancers designed by in silico sequence evolution or motifs implantation. Nevertheless, some of the most active GAN-generated sequences in the MEL line (e.g. GG-3 and GG-4) are highly specific, as they have very low activity in this MES line (Supplementary Fig. 8b).

In conclusion, these results show that enhancer design strategies are adaptable to other species including human, and to different biological systems.

## Discussion

Understanding the code of enhancers and utilising this knowledge to design synthetic enhancers is a long-standing problem. Here we designed synthetic enhancer sequences in human and fly guided by deep learning models. We used models that we previously trained and validated on cell type or cell state-specific chromatin accessibility ^11,44,49^. All these models are freely available from the Kipoi repository ^54^, and can be re-used by the community to design enhancers. During the enhancer design process, we gained new insight into enhancer codes: an enhancer sequence can emerge from a random or genomic sequence through a relatively small number of mutations; a second code can be added to an enhancer to make it active in an additional cell type; intertwined enhancer codes can be untwined, so that its activity is pruned to a specific cell type; enhancers can be minimised to the extreme situation where they only contain TF motifs; and generative models can deliver functional and cell type specific enhancers.

By combining a stepwise enhancer design approach alongside model interpretation techniques, we followed the trajectories of in silico enhancer emergence in Drosophila and human. Nucleotide-by-nucleotide evolution revealed that the selected mutations predominantly destroy repressor TF binding sites and create activator sites. A single mutation can also convert a repressor binding site directly into an activator site and such mutations were most preferred. In most cases, ten iterative mutations were sufficient to convert a random sequence into a cell type specific functional enhancer. These findings are corroborated by a recent study where it was shown that the expression of native yeast promoter sequences can increase to extremely high or low levels with only four mutations ^45^. This evolutionary design process may also reflect natural evolution of genomic enhancers. Indeed, we found that the fly genome contains many “near-enhancers” that require few mutations to become a functional enhancer. This finding highlights the potential of these sequences to become enhancers, or alternatively reflect evolutionary remnants.

Even though we directed the sequence evolution by following the prediction score towards a single cell type, without taking other cell types into account, the generated sequences were mostly active only in the targeted cell type. This suggests that: (1) activator binding sites were not created for other cell types; and (2) repressor sites, which are present in random sequences by chance, were not destroyed for other cell types. For example, in Kenyon Cells we observed that activator binding sites are usually longer than repressor sites (18 bp and 10 bp versus 5 bp and 6 bp for Ey, Mef2, Mamo, and CAATTA respectively). This implies that a random sequence is more likely to have multiple repressor binding sites by chance compared to activator sites. Indeed, the average prediction scores of our initial 6,000 random sequences were close to zero for all classes. This may at least in part explain why earlier enhancer design efforts may have failed ^55^ : when TF motifs are implanted at random positions in a random sequence, repressor sites may remain present. Likewise, to be able to generate a functional enhancer through random sequencing generation, many random sequences need to be generated (e.g., 100 million and 1 billion ^42,56^).

The location, orientation, strength, and number of TF motifs within a single enhancer, and its distance to other motifs, is what defines a unique enhancer code. This array of well-positioned TF binding sites constitutes a possible docking platform for a specific combination of TFs. Their cooperative binding makes the enhancer accessible/active at different levels and in different cell types. Under this premise, how can a single enhancer be active in multiple cell types, or at multiple time points? Besides the trivial possibility whereby the two cell types share a common set of TFs that bind to a common set of sites (e.g., different KC subtypes), we discovered here that some enhancers have evolved with multiple intertwined codes (e.g., KCs and T neurons). We could prove this by either removing a code from a native dual-code enhancer, or adding a second code to a native single-code enhancer. In other words, a dual-code enhancer can be mutated in such a way that only one of the codes is affected.

Design by in silico evolution suggests that enhancers consist merely of correctly positioned motifs, which is in agreement with the cooperative enhancer model, with flexible motif syntax ^2,41^. The various enhancer models we studied here seem less ‘tightly fixed’ compared to the enhanceosome model, but more constrained than the ‘bag of motifs’ or billboard model ^57^. Indeed, melanoma enhancers remained active when motifs were moved to less preferred locations. The consequence of this motif-dictated enhancer model is that it allows for enhancer design by motif implantation. Several studies have used motif implantation in an attempt to design enhancers, but successes of accurate in vivo activity have been limited ^55^. Deep learning models provide the advantage that many different motif implanting scenarios can be tested in silico, before performing any experimental validation ^41,42^, compared to high-throughput testing of random implantations ^31,58,59^. By exploiting this technique further, particularly by scoring each possible implant position, as well as combinations of motifs, we could reveal motif synergies (e.g., Ey + Mef2; or SOX10 + MITF), as well as preferred orientations and distances between motifs, motif strengths, and motif copy number. In an extreme case, a minimal fly brain enhancer could be designed with three abutting motif instances, illustrating that functional enhancers can be created without any further sequence context.

An entirely different type of enhancer design uses GANs. GANs have proven their strengths in the generation of realistic samples in many different fields and they have been proposed to generate enhancer sequences *in silico* ^51^. In our experience, by only using around 4000-6000 genomic sequences as training samples, GAN models successfully generated sequences of which around 30-50% were correctly predicted to be specific for the trained class. This might be further improved in the future by using a larger set of real samples (e.g., cross-species and multiple cell types) or by improving network architectures. Even though GANs could generate functional enhancers, we needed a trained deep learning model to measure the quality of the generated sequences. Indeed, to our knowledge there is not a simple technique to measure the performance of the generator and the discriminator during training. In this aspect, sequence evolution and motif implantation approaches were more interpretable and straightforward compared to generative design.

The successful application of enhancer design strategies guided by deep learning models on both fly brain and human cancer cells have shown that these simple (yet very powerful) strategies are adaptable to any organism or system. Our proof-of-concept study is an encouraging step forward towards the development of organism-wide deep learning models. Such models will facilitate the generation of synthetic enhancers during development, disease, and homeostasis; and will further improve our understanding and control of the genomic cis-regulatory code.

## Methods

### In silico saturation mutagenesis

To measure the effect of each possible single mutation on a given DNA sequence, we performed in silico saturation mutagenesis. We first generated the sequences of all single mutations for a given 500 bp sequence (3 possible mutations for each nucleotide, making 1500 sequences in total). We scored these sequences and the initial sequence with the deep learning models. For a chosen class, we calculated the delta prediction score by subtracting the score of the initial sequence from the score of the mutated sequence for each mutation.

### Random sequence generation

We generated random 500 bp sequences to use as a prior set for the in silico sequence evolution and motif implantation by using the *numpy*.*random*.*choice([“A”,”C”,”G”,”T”])* command. For each position, instead of using 25% probability for each nucleotide to be chosen, we used the frequency of the nucleotides from fly and human genomic regions for each position. In these genomic regions, the GC-content was higher in the center of the regions on average relative to the flankings. We used 6126 KC regions for fly and 3885 MEL regions for human that we identified in our previous publications ^44,49^.

### In silico sequence evolution

By using the saturation mutagenesis scores mentioned above, we performed in silico sequence evolution. For the in silico evolution from random sequences, we calculated saturation mutagenesis scores for a random sequence. Then, we selected the mutation that had the highest positive delta prediction score for the selected class (for γ-KC, class no. 35 in DeepFlyBrain; for MEL, class no. 16 in DeepMEL2). For the selected sequence with one mutation, we recalculated the saturation mutagenesis scores for each nucleotide and again selected the mutation with the highest delta score and repeated this procedure until the initial random sequence accumulated 20 mutations. We used 6000 initial random sequences for γ-KC and 4000 for MEL. For the generation of KC enhancers from genomic regions, we performed 6 iterative mutations. For the multiple cell-type code enhancers, we started from optic lobe enhancers and in each iteration we selected the mutations that increased the γ-KC prediction score while maintaining the optic lobe prediction scores high. For the pruning experiment of a multiple cell type code enhancer into only KC code, we selected the mutations that maintain the γ-KC prediction score high while decreasing the optic lobe prediction scores. The DeepFlyBrain class numbers used for optic lobe neurons are 23 for T1, 20 for T2, and 2 for T4 neurons.

### Nucleotide contribution scores

We used a network explaining tool, called DeepExplainer (SHAP package ^47^), to calculate the contribution of each nucleotide to the final prediction of the deep learning model for the chosen class. We used randomly selected 250 genomic regions to initialize the explainer.

DeepFlyBrain model takes a single strand as an input. For a given 500 bp, we multiplied the explainer’s output by the one-hot encoded DNA sequence and visualized it as the height of the nucleotide letters. DeepMEL2 model takes forward and reverse strands separately as an input. In this case, the explainer results in contribution scores for each strand. We first took the average contribution score for each nucleotide and then multiplied it by the one-hot encoded DNA sequence to visualize.

### Scoring the fly genome

To identify the regions that have high prediction scores for γ-KC but have less accessibility in γ-KC, we scored the whole fly genome. We used the *bedtools makewindows -g dm6*.*chromsize -w 500 -s 50* command ^60^ to create the coordinates of the binned fly genome with a 500 bp window and 50 bp stride. We removed the regions that are not exactly 500 bp. This resulted in 2,750,893 regions to be scored with the DeepFlyBrain model. We used the *stats* function of deeptools/pyBigWig package ^61^ to calculate mean γ-KC accessibility values for each bin.

### Motif implanting

To implant binding sites into 500 bp sequences, we started from a random sequence. We implanted a binding site into every possible location on the random sequence one-by-one by replacing the nucleotides on the random sequences with the binding site. Then, we scored these sequences with the model. We selected the binding site position that gives the highest prediction score and implanted the motif on that position. Then, starting from this sequence with one binding site implanted, we implanted the next binding sites one-by-one by using the same procedure. The sequence of implanted binding sites are as follows; Ey: TGCTCACTCAAGCGTAA, Mef2: CTATTTATAG, Onecut: ATCGAT, Sr: CCACCC, SOX10: AACAATGGGCCCATTGTT, MITF: GTCACGTGAC, and TFAP2: GCCTGAGGC. We used 2000 initial random sequences for γ-KC and 2000 for MEL. The weaker binding sites taken from the IRF4 enhancer are as follows: SOX10_1: GTGAATGACAGCTTTGTT, SOX10_2: TACAAGTATCTCCATTGT, MITF_1: ATCATGTGAA, MITF_2: GCCATATGAC, TFAP2_1: TCTTCAGGC, and TFAP2_2: CCCTGTGGT.

### Generative Adversarial Network

To train a GAN model, we used Wasserstein GAN architecture with gradient penalty ^46^ similar to earlier work ^51^. The model consists of two parts; generator and discriminator. Generator takes noise as input (size is 128), followed by a dense layer with 64,000 (500 * 128) units with ELU activation, a reshape layer (500, 128), a convolution tower of 5 convolution blocks with skip connections, a 1D convolution layer with 4 filters with kernel width 1, and finally a SOFTMAX activation layer. The output of the generator is a 500 × 4 matrix, which represents one-hot encoded DNA sequence. Discriminator takes 500 bp one-hot encoded DNA sequence as input (real or fake), followed by a 1D convolution layer with 128 filters with kernel width 1, a convolution tower of 5 convolution blocks with skip connections, a flatten layer, and finally a dense layer with 1 unit.

Each block in the convolution tower consists of a RELU activation layer followed by 1D convolution with 128 filters with kernel width 5. The noise is generated by the *numpy*.*random*.*normal(0, 1, (batch_size, 128))* command. We used a batch size of 128. For every *train_on_batch* iteration of the generator, we performed 10 *train_on_batch* iteration for the discriminator. We used Adam optimizer with learning_rate of 0.0001, beta_1 of 0.5, and beta_2 of 0.9. We trained the models for around 260,000 batch training iteration for KC and around 160,000 batch training iteration for MEL.

We used 6,126 KC regions for the fly model and 3,885 MEL regions for the human model, which we identified in our previous publications, as real genomic sequences to train the models. After the training, we sampled 6,144 (48 * batch size) sequences for KC and 3,968 (31 * batch size) sequences for MEL by using the generator for every 10,000 batch training iteration. The sampled synthetic sequences were generated by calculating predictions on noise and then the *numpy*.*argmax()* command was used to convert the predictions into one-hot encoded representations.

### Background model

To compare against the GAN-generated sequences, we generated random sequences in different orders by using the *CreateBackgroundModel* function from the INCLUSive package ^62^ based on the same genomic regions that we used to train GANs.

### Cloning of synthetic Drosophila enhancers

Synthetic sequences (except for the motif-implantation and double-coded sequences) were ordered from Twist Bioscience, pre-cloned in the pTwist ENTR vector. The motif-implantation and double-coded sequences were synthesized with an additional 5’ CACC sequence as double-stranded DNA (gBlocks Gene Fragments) by IDT. 49 bp motif-implantation sequence was ordered from IDT as forward and reverse single-stranded DNA oligos, which were then annealed for 5 min at 95°C and cooling down to RT over one hour. The double-stranded DNA sequences were then cloned into the pENTR/D-TOPO plasmid (Invitrogen).

All sequences were introduced in the pH-Stinger vector, containing nuclear GFP, Hsp70 promoter and gypsy insulators, via Gateway LR recombination reaction (Invitrogen) and 2 µl of the reaction was transformed into 25 µl of Stellar chemically competent bacteria (Takara). Plasmid minipreps were performed using the NucleoSpin Plasmid Transfection-grade Mini kit (Macherey-Nagel) and sequenced with Sanger sequencing to confirm the correct insertion of the regions in the destination plasmid. Next, the plasmids were sent to FlyORF (CH) for injection in Drosophila embryos (21F site on chromosome 2l) and positive transformants were selected.

Drosophila flies were raised on a yeast-based medium at 25°C under a 12 h-12 h day-night light cycle.

### Immunohistochemistry analysis of Drosophila brains

Brains of adult flies (<10 days old) were dissected in PBS and transferred to a tube for fixation in 4% formaldehyde in PBS for 20 min. All incubations were done at room temperature, unless otherwise indicated. Brains were washed in PBS with 0.3% Triton-X (PBST) three times for 10 min each, then they were placed in blocking solution (5% normal goat serum [Abcam] in PBST) for 3 hours. We incubated the brains overnight at 4°C in primary antibody (rabbit anti-GFP, IgG [Invitrogen]), diluted 1:1000 in blocking solution. The brains were then washed in PBST three times for 10 min each and then incubated with the fluorochrome-conjugated secondary antibody Alexa Fluor 488 donkey anti-rabbit IgG (Invitrogen), diluted 1:500 in blocking solution for 2 hours. Next, brains were washed in PBS three times for 10 min each. Finally, samples were mounted onto microscope slides with Prolong Glass Antifade Mountant (Invitrogen).

For image acquisition, a Zeiss LSM900 microscope equipped with Airyscan2 in combination with a 20x objective (Plan Apo 0,80 Air) was used. The setup was controlled by ZEN blue (version 3.4.91, Carl Zeiss Microscopy GmbH). GFP was excited with a blue diode 100mW at 488 nm and tiled images were collected with emission filter BP450-490/BS495/BP500-550.

### Cloning of synthetic human enhancers

500 bp synthetic sequences were ordered from Twist Bioscience, pre-cloned in the pTwist ENTR vector. 500 bp regions were introduced in the pGL4.23-GW luciferase reporter vector (Promega) via Gateway LR recombination reaction (Invitrogen) and 2 µl of the reaction was transformed into 25 µl of Stellar chemically competent bacteria (Takara).

Synthetic sequences shorter than 150 bp were ordered as gBlocks from IDT (Integrated DNA Technologies) with 5’ (cccgtcgacgaattctgcagatatcacaagtttgtacaaaaaagcaggct) and 3’ (acccagctttcttgtacaaagtggtgataaacccgctgatcag) adaptors. The pGL4.23-GW luciferase reporter vector was linearized via inverse PCR with primers Lin_pSA335_short_ME_For (gtggtgataaacccgctgatcag) and Lin_pSA335_short_ME_Rev (tctgcagaattcgtcgacggg). The short sequences and the linearized vector were combined in an NEBuilder reaction (New England Biolabs, Ipswich, MA) and 2 µl of the reaction was transformed into 25 µl of Stellar chemically competent bacteria.

For all cloning procedures, plasmid minipreps were performed using the NucleoSpin Plasmid Transfection-grade Mini kit (Macherey-Nagel) and sequenced with Sanger sequencing to confirm the correct insertion of the regions in the destination plasmid.

### Transfection and luciferase assay

MM001 and MM047 were seeded in 24-well plates and transfected with 400 ng pGL4.23-enhancer vector + 40 ng pRL-TK renilla vector (Promega) with Lipofectamine 2000 (Thermo Fisher Scientific). As positive controls, the previously published enhancers MLANA_5-I, IRF4_4-I and TYR_-9-D or ABCC3_11-I and GPR39_23-I were used for MM001 and MM047 respectively ^53^. One day after transfection, luciferase activity was measured via the Dual-Luciferase Reporter Assay System (Promega) by following the manufacturer’s protocol. Briefly, cells were lysed with 100 µl of Passive Lysis Buffer for 15 min at 500 rpm. 20 µl of the lysate was transferred in duplicate in a well of an OptiPlate-96 HB (PerkinElmer, Waltham, MA) and 100 µl of Luciferase Assay Reagent II was added in each well. Luciferase-generated luminescence was measured on a Victor X luminometer (PerkinElmer). 100 µl of the Stop & Glo Reagent was added to each well, and the luminescence was measured again in order to record renilla activity. Luciferase activity was estimated by calculating the ratio luciferase/renilla; This value was normalized by the ratio calculated on blank wells containing only reagents. Three biological replicates were done per condition for MM001 and two biological replicates for MM047.

## Data availability

Cloned Drosophila and human sequences were provided as Supplementary Tables. DeepMEL, DeepMEL2, and DeepFlyBrain deep learning models were obtained from Kipoi ^54^ (http://kipoi.org/models/DeepMEL, https://kipoi.org/models/DeepFlyBrain). The fasta files used to train GAN models and the trained GAN models are available on Zenodo at https://doi.org/10.5281/zenodo.6701504. Chromatin accessibility values in Kenyon Cells in adult Drosophila brains were obtained from GSE163697 ^49^.

## Acknowledgements

The authors gratefully acknowledge the VIB Bio Imaging Core for their support & assistance in imaging. Computing was performed at the Vlaams Supercomputer Center (VSC). This work is funded by the following grants to S. Aerts: ERC Consolidator Grant (724226_cis-CONTROL), ERC Proof of Concept (963884), Special Research Fund (BOF) KU Leuven (grant C14/18/092), Foundation Against Cancer (2020-062), and FWO (grant G094121N).

## Author contributions

I.I.T. and S.A. conceived the study; I.I.T. performed all computational analyses and designed synthetic enhancers; V.C. performed enhancer cloning experiments with assistance from K.I.S and D.M.; V.C. performed luciferase assays with assistance from D.M.; K.I.S. performed antibody staining and visualization with assistance from I.I.T.; I.I.T. and S.A. wrote the manuscript with assistance from D.M..

## Supplementary Figures

**Supplementary Figure 1:**
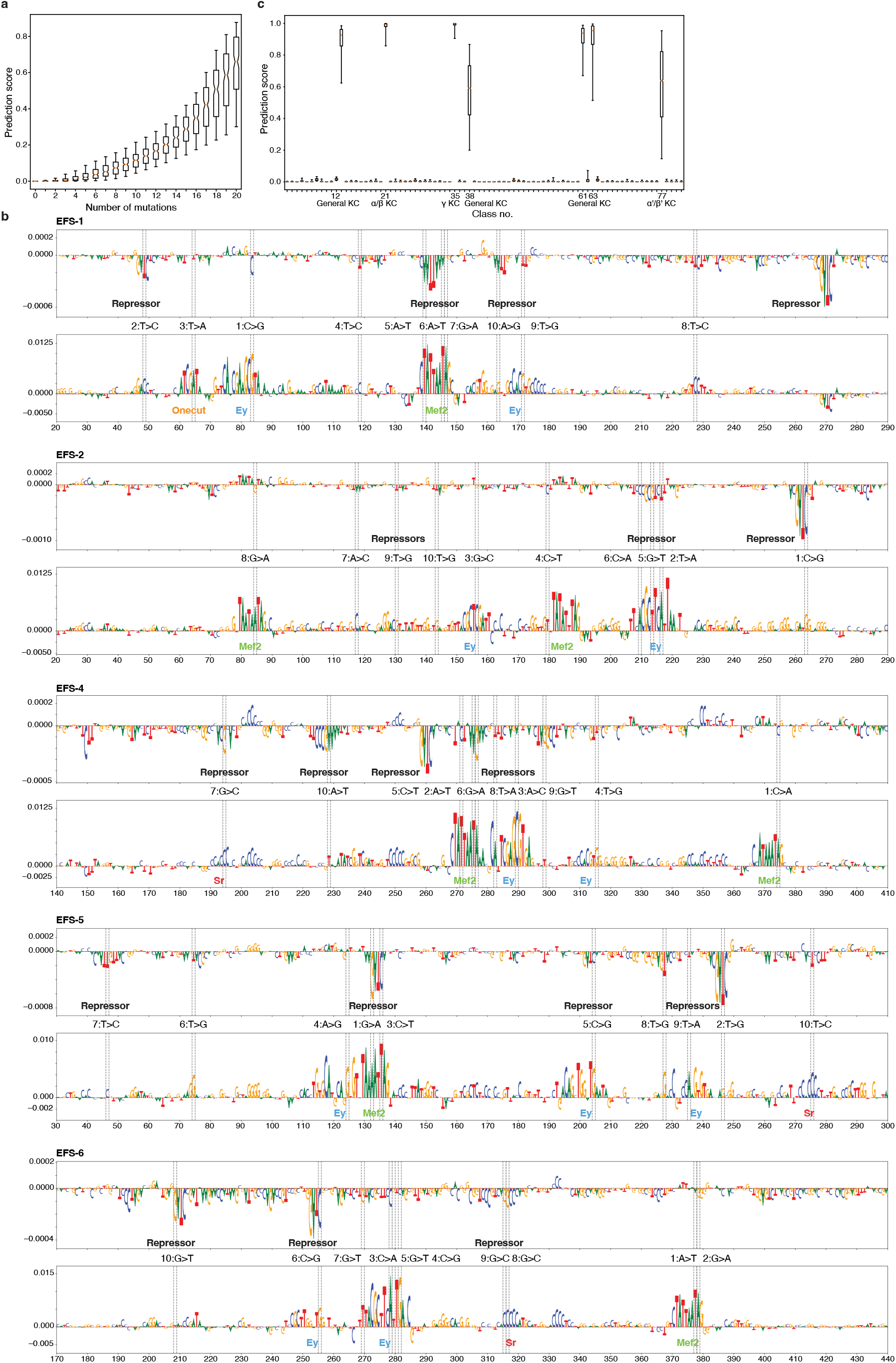
In silico sequence evolution from random sequencesa. **a**, Prediction score distribution of the sequences that do not reach 0.5 prediction score threshold after 15 mutations for the γ-KC class (*n* = 180) after each mutation. **b**, Nucleotide contribution scores of selected random sequences in their initial form (top) and after 10 iterations (bottom). The position of the mutations are shown with dashed lines. The mutational order is written in between top and bottom plots together with the type of nucleotide substitutions. **c**, Prediction score distribution of selected regions (*n* = 6) for all classes after 10 mutations. The KC specific classes and their class number are indicated. In **a, c**, the box plots show the median (center line), interquartile range (box limits), and 5th and 95th percentile range (whiskers).

**Supplementary Figure 2:**
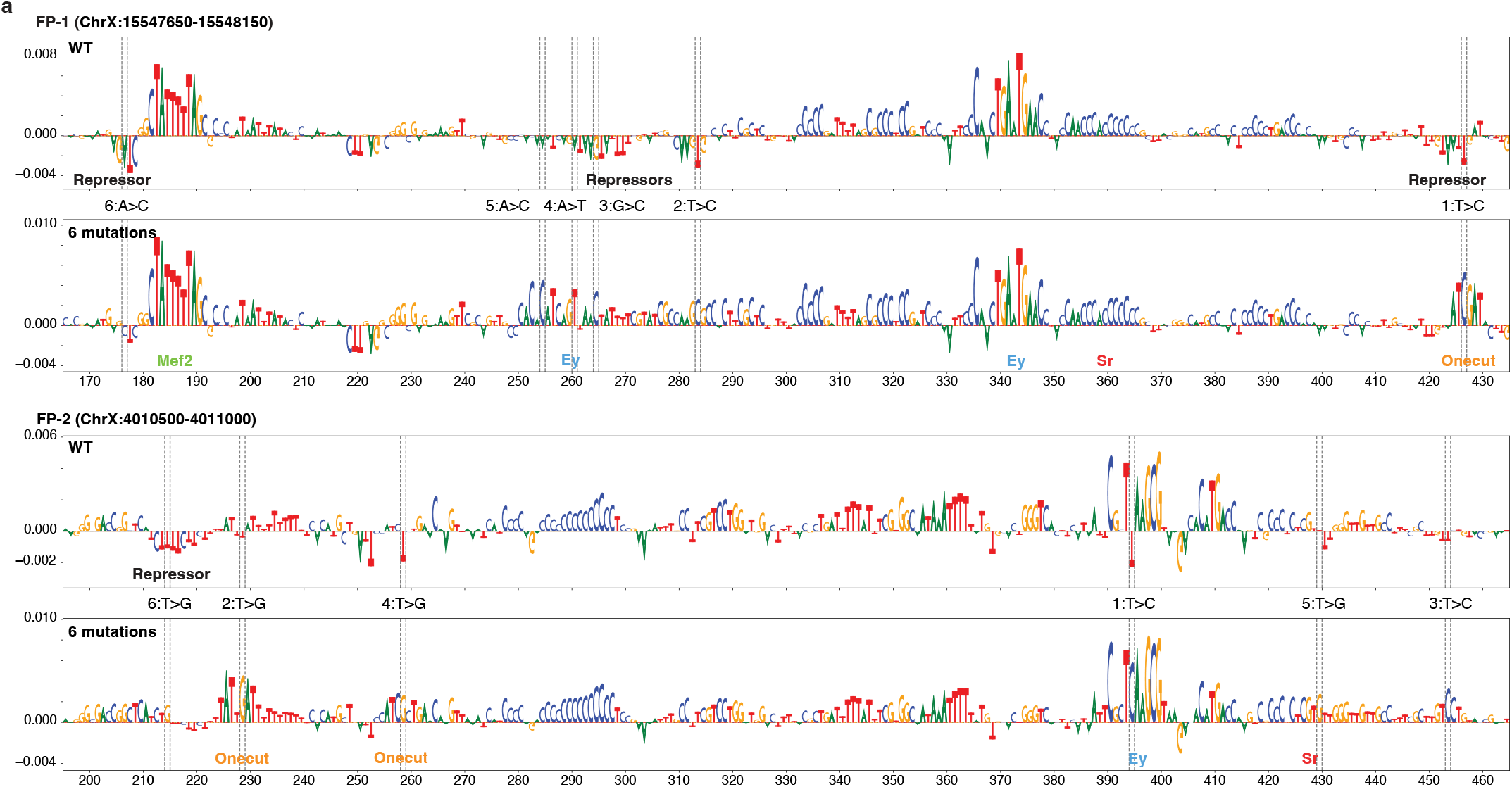
In silico sequence evolution from genomic sequences. **a**, Nucleotide contribution scores of selected genomic sequences in their initial form (top) and after 6 iterations (bottom). The position of the mutations are shown with dashed lines. The mutational order is written in-between top and bottom plots together with the type of nucleotide substitutions.

**Supplementary Figure 3:**
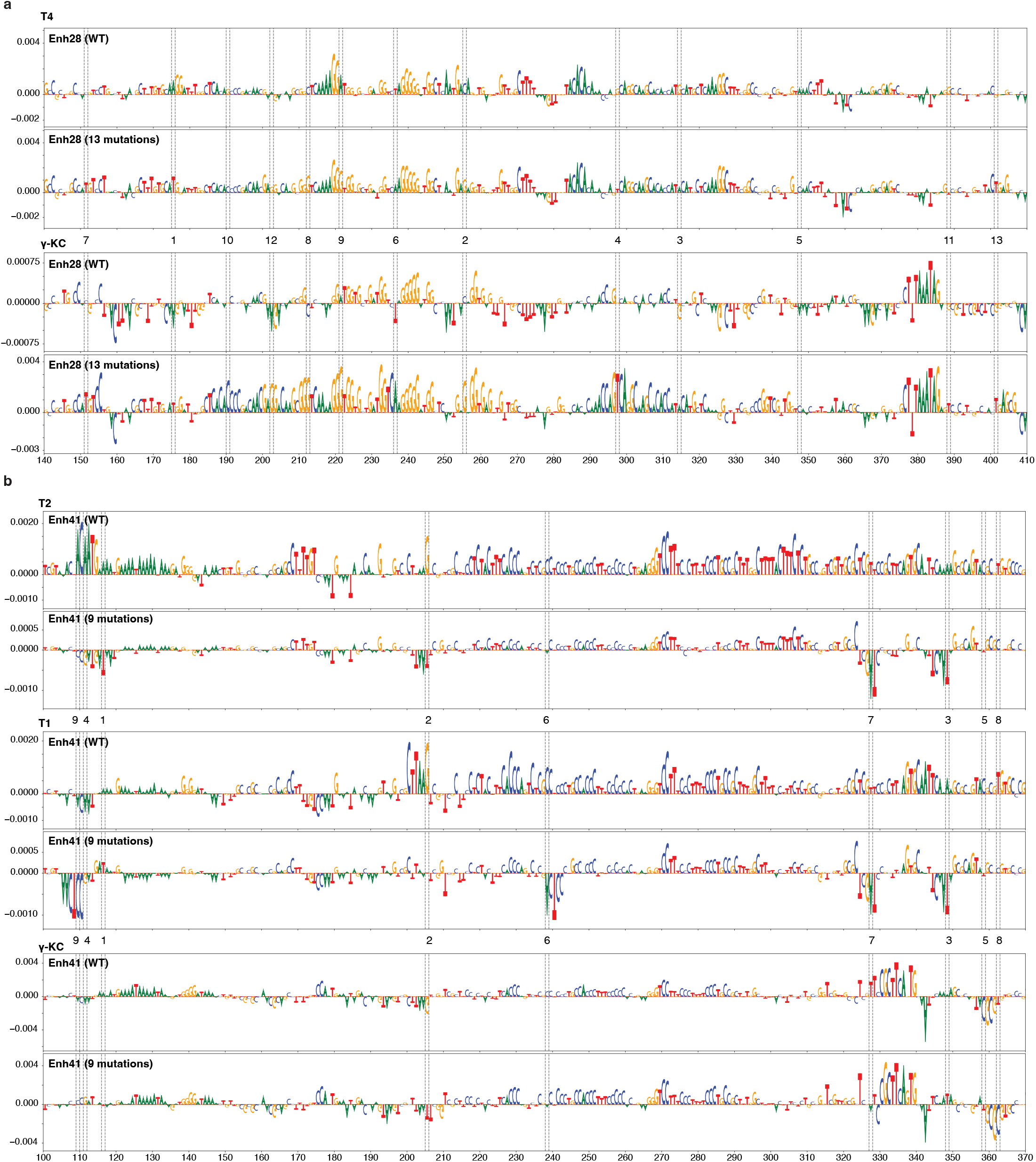
Enhancer design towards multiple cell type codes. **a**, Nucleotide contribution scores of wild-type (WT) sequence of the *amon* enhancer and after 13 mutations for T4 (top) and γ-KC (bottom). The position of the mutations are shown with dashed lines. The mutational order is written in-between top and bottom plots. **b**, Nucleotide contribution scores of wild-type (WT) sequence of the *Pkc53e* enhancer and after 9 mutations for T2 (top), T1 (middle), and γ-KC (bottom). The position of the mutations are shown with dashed lines. The mutational order is written in-between top and bottom plots.

**Supplementary Figure 4:**
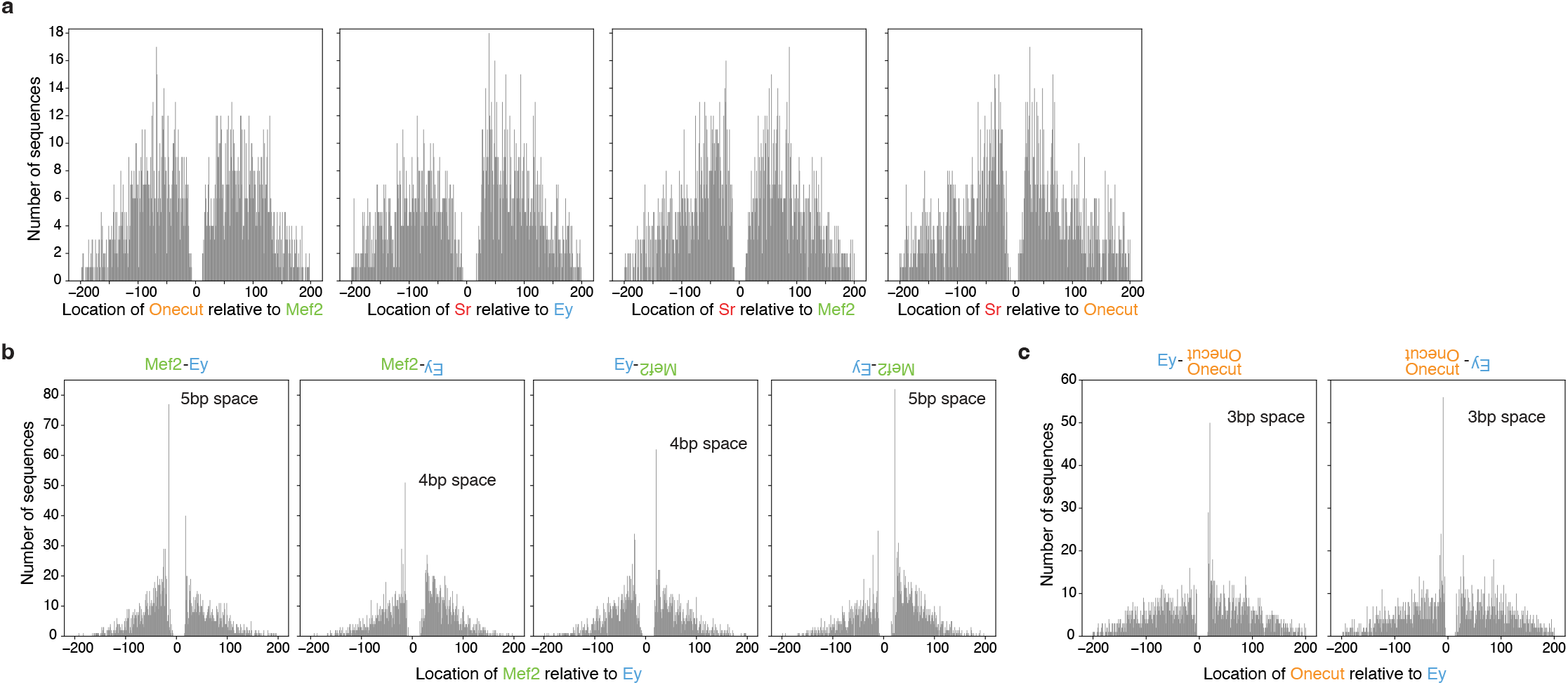
Preferred distances between implanted binding sites. **a**, Distribution of Onecut locations relative to Mef2, Sr to Ey, Sr to Mef2, and Sr to Onecut, respectively (*n* = 2,000). **b**, Distribution of Mef2 locations relative to Ey when both are on the same strand, Ey is on the negative strand, Mef2 is on the negative strand, and both are on the negative strand, respectively (*n* = 2,000). **c**, Distribution of Onecut locations relative to Ey when Ey is on the positive strand and when Ey is on the negative strand, respectively (*n* = 2,000).

**Supplementary Figure 5:**
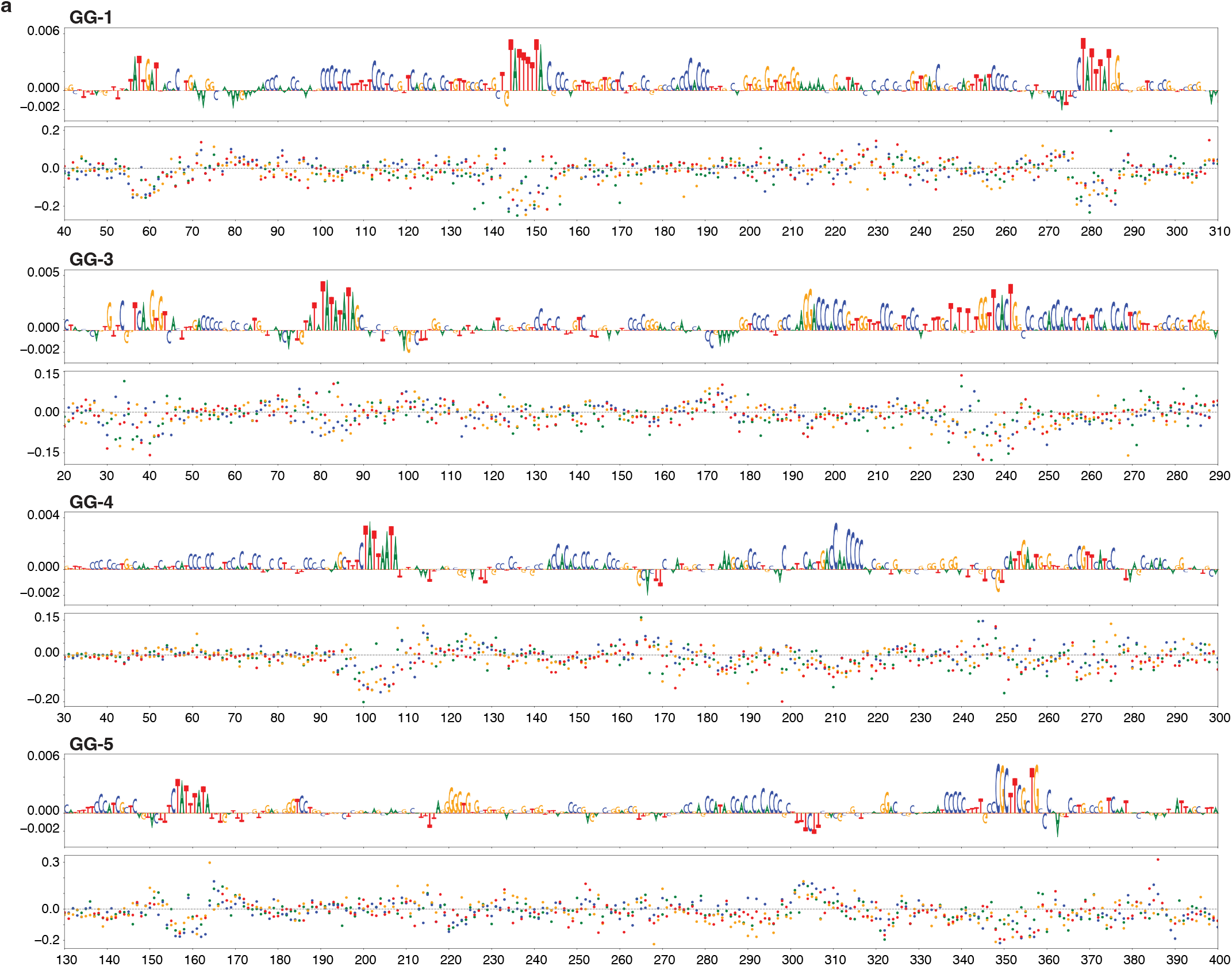
Generative design of enhancers. **a**, Nucleotide contribution scores of selected GAN-generated sequences (top) and their in silico saturation mutagenesis assays (bottom). Each dot on the saturation mutagenesis plot represents a single mutation and its effect on the prediction score (*y* axis).

**Supplementary Figure 6:**
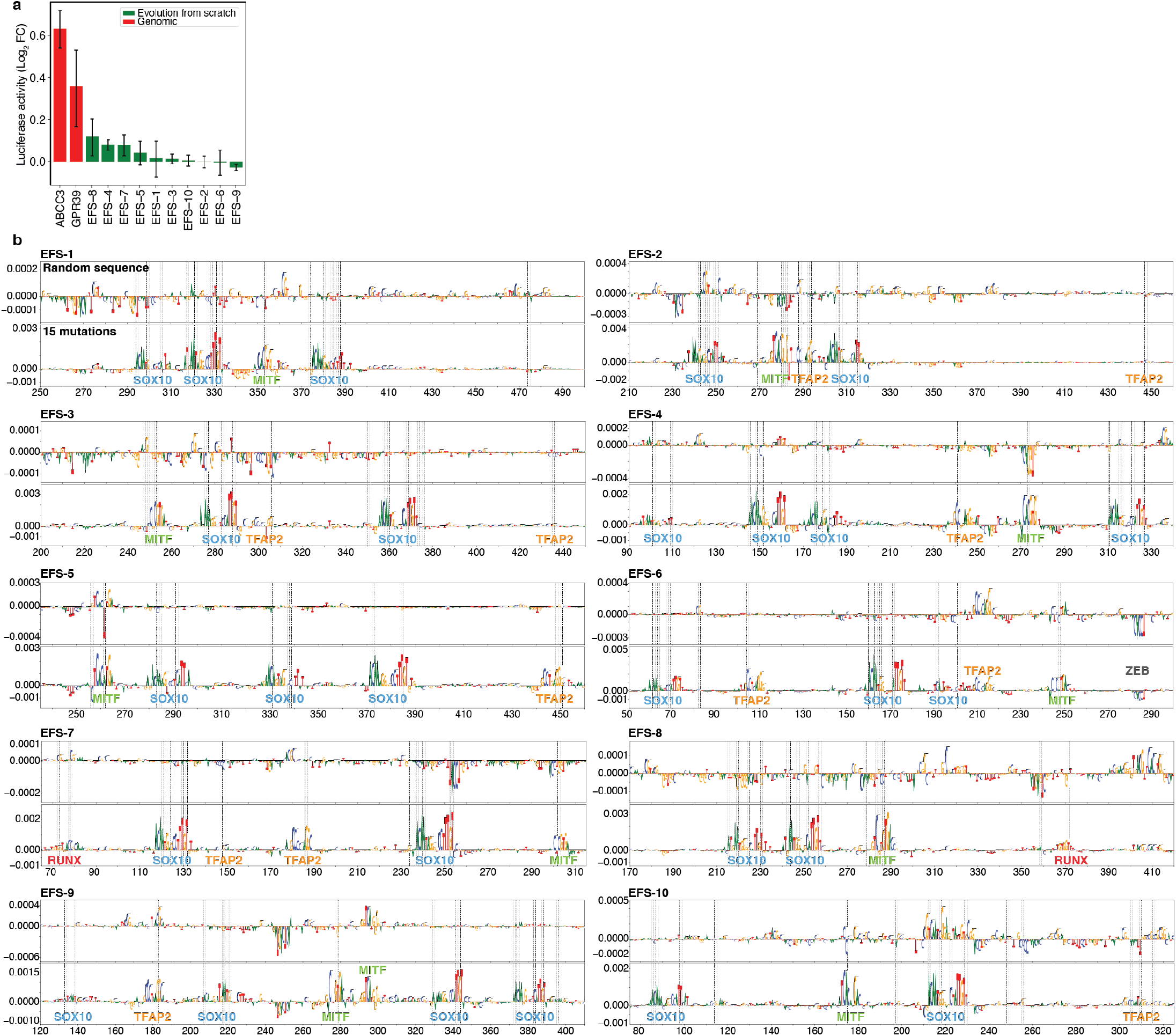
Human enhancer design by in silico evolution. **a**, Bar plot showing the mean luciferase signal (log_2_ fold-change over Renilla) in a MES melanoma line (MM047) of the synthetic MEL enhancers (generated by in silico sequence evolution), showing no activity compare to positive control genomic MES enhancers. The error bars show the standard error of the mean (*n* = 2 biological replicates). **b**, Nucleotide contribution scores of the selected synthetic sequences in their initial form (top) and after 15 iterations (bottom). The position of the mutations are shown with dashed lines. The name of the generated binding sites by the mutations are indicated at the bottom.

**Supplementary Figure 7:**
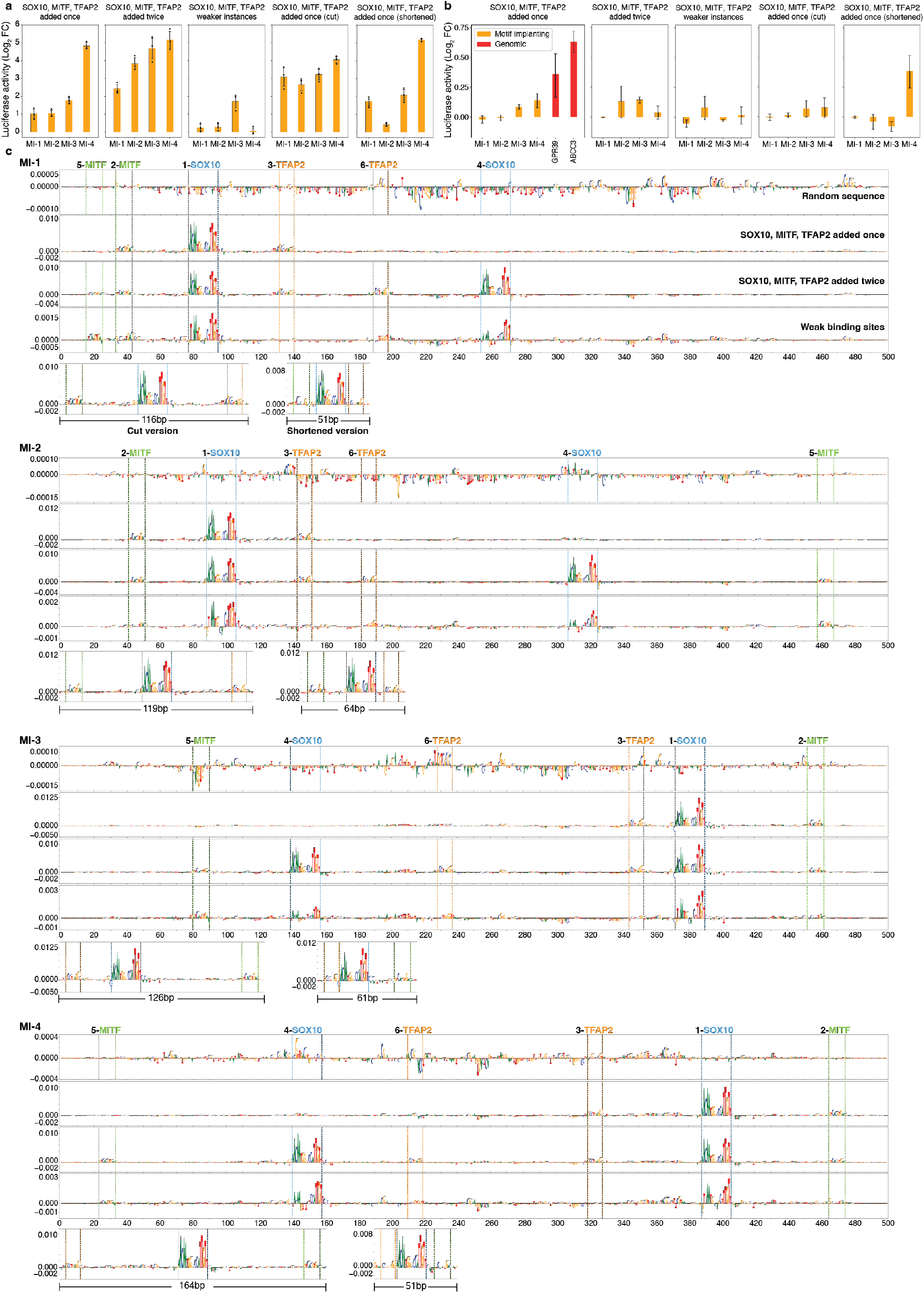
Human enhancer design by motif implantation. **a-b**, Bar plots show the mean luciferase signal (log_2_ fold-change over Renilla) of the synthetic sequences, which were generated by motif implantation, tested in MM001 (**a**, MEL melanoma cell line, *n* = 3 biological replicates) and MM047 (**b**, MES melanoma cell line, *n* = 2 biological replicates). Values of 2 previously validated MES regions are displayed for MM047. The error bars show the standard error of the mean. **c**, Nucleotide contribution scores of the selected synthetic sequences in their initial form (first row), after adding SOX10, MITF, and TFAP2 motifs once (second row), after adding SOX10, MITF, and TFAP2 motifs twice (third row), weaker-motif version of the third row after replacing implanted motifs with weaker sites (fourth row), cut version of the second row where only the part with the binding sites were taken (fifth row, left), and shortened version of the second row where MITF and TFAP2 placed as close as possible to SOX10 (fifth row, right). The name of the motifs and their implantation order are indicated at the top. The position of the motifs are shown with dashed lines.

**Supplementary Figure 8:**
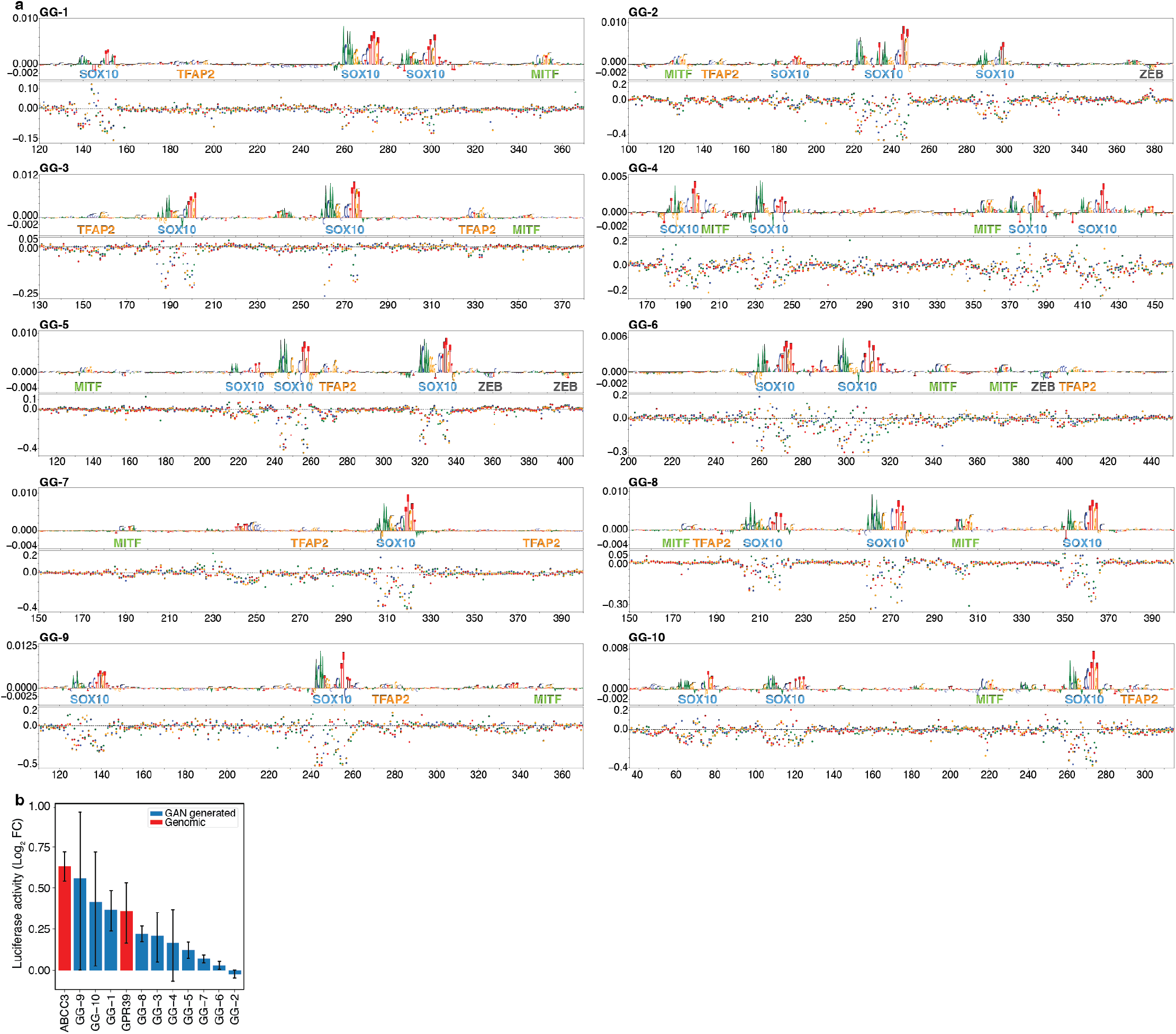
Human enhancer design by Generative Adversarial Networks. **a**, Nucleotide contribution scores of the selected GAN-generated sequences (top) and their in silico saturation mutagenesis assays (bottom). Each dot on the saturation mutagenesis plots represents a single mutation and its effect on the prediction score (*y* axis). The name of the identified binding sites are written in between top and bottom plots. **b**, Bar plot showing the mean luciferase signal (log_2_ fold-change over Renilla) of the GAN-generated sequences and genomic MES enhancers in MM047 (MES melanoma line). The error bars show the standard error of the mean (*n* = 2 biological replicates).

